# Microparticle-Enabled Single Cell Multiparameter Electronic Immunophenotyping for Selective Electroporation

**DOI:** 10.1101/2025.10.17.682894

**Authors:** Madeline Hoyle, Josiah Rudge, Yuvraj Rallipalli, Aniruddh Sarkar

## Abstract

Electroporation (EP) is one of the leading non-viral intracellular delivery methods used in various applications across research and cell therapy development and manufacturing. Currently widely used bulk EP methods, while they offer scalability, cost efficiency and simplicity, cannot be used for targeted or selective delivery to a defined subset of a input cell population. Here, we present a Microparticle-Enabled Selectively Permeabilizing Impedance Cytometer (ME-SPICy), a microfluidic single-cell EP platform that enables targeted EP of selected cell subpopulations based on their surface markers. Antibody conjugated microparticles (MPs) are used to label selected cell subpopulations within a larger heterogenous sample. Using multifrequency impedance detection, ME-SPICy discriminates, in real-time, non-labeled and labeled cells within the mixed sample as they flow through a 3D printed biconical micro-aperture. This allows for the system to analyze if a cell is a target cell and selectively apply a low voltage (<16 V) for targeted single-cell EP. Simulations and experimental validation demonstrate that MP binding substantially alters cell impedance and phase signature, enabling accurate label-based discrimination. We demonstrated selective EP first using Jurkat cells by targeting either the labeled or non-labeled populations. Then we demonstrated targeted delivery to primary human lymphocytes within peripheral blood mononuclear cells. ME-SPICy achieved high precision, with 98% purity and >5 fold enrichment of lymphocytes in the electroporated cell population. This approach expands the capabilities of EP, offering a promising solution to decrease manufacturing complexity in both research and clinical cell engineering workflows

## Introduction

Engineered cell therapies have emerged as a promising treatment option for many cancers and diseases, with thousands of clinical trials either completed or in progress[1]. A critical step in engineering, or modifying, cells for these therapies is the intracellular delivery of gene editing cargo across the cell membrane. This delivery can be achieved by using chemical, biological, or physical methods[2]. Chemical methods, such as lipofection, while commonly used in research settings, can result in inconsistent delivery, especially with primary cells [3-5]. Viral vector-based methods are popular due to their high efficiency, often exceeding 90%, but can have significant limitations too [6, 7]. These include high costs, lengthy production time for clinical grade vectors, limited cargo payloads, and lingering long-term safety concerns[8][6].

Physical methods to permeate the cell membrane, such as electroporation (EP), address many of the challenges associated with viral methods [9]. In conventional EP for intracellular delivery, a suspension of cells and deliverable cargo is placed between two electrodes, typically spaced a few millimeters apart. A high-voltage pulse (>1000 V) is applied across these electrodes. This creates an electric field which increases the transmembrane potential (TMP) of the cells, inducing transient pores in the cell membrane and allowing cargo to enter [10, 11]. However, conventional EP methods can experience inconsistent delivery efficiency and reduced post-EP cell viability, especially for heterogeneous primary cells, due to electrolysis and Joule heating from the high voltages required [11, 12].

Microfluidic EP systems have emerged as a subtype of electroporation that operate at significantly lower voltages due to their microscale architecture. These systems offer improved efficiency and viability by minimizing electrolysis [4, 12-16]. Additionally, they allow for single-cell resolution and can be integrated with fluorescent imaging or electrical measurements for real-time monitoring or precise control of EP parameters [17-22]. Multifrequency impedance cytometry, an electrical measurement technique, enables label-free single-cell phenotyping based on cellular dielectric properties [23-26]. At low measurement frequencies (<1 MHz), the impedance signal correlates with cell size and volume, while at higher frequencies (>1 MHz), it is related to membrane and intracellular properties [24]. However, most current biological and clinical cell targeting needs depend on cell identification via differential expression of cell surface markers. Microparticles have shown promise in their use as tags to convert surface marker expression to a size or impedance signature for affinity-based multiplexed cell sorting [27] and impedance characterization as well [25].

When combined with single-cell microfluidic EP, multifrequency impedance cytometry has the potential to identify and selectively electroporate subpopulations of cells, enabling targeted EP which is not achievable with conventional EP methods. Here, we present a Microparticle Enabled Selectively Permeabilizing Impedance Cytometer (ME-SPICy) to achieve selective EP of a cell subpopulation within a heterogenous mixture. Cells are flowed through a 3D printed microchannel where we use multi-frequency impedance cytometry to identify single cells and determine, in real time, if a cell is labeled with an MP and thus is a target cell for electroporation. Once a target cell is identified, a low voltage DC pulse is applied to electroporate the cell. This enabled selective EP within cell lines and primary cells for both labeled and non-labeled populations. This method expands the capabilities of EP while also being a cost-effective and efficient alternative to viral vectors for targeted intracellular delivery.

## Results and Discussion

### Principle and Microfluidic System

A 3D printed biconical vertical micro-aperture is coupled with fluidics and electronics for multifrequency impedance measurements [28] and real-time discrimination of non-labeled and MP-labeled cells for selective EP. Cells flow rapidly through the narrowing micro-aperture which concentrates the electric field at a constriction (Figure 1a). This geometry ensures that only one cell passes through the constriction at a time. Additionally, at the narrowest part of the constriction, the cell experiences a high electric field, as evidenced by COMSOL simulations (Figure 1b) while using a much lower voltage (∼15V) compared to bulk EP (∼100V-1000V).

**Figure 1.**
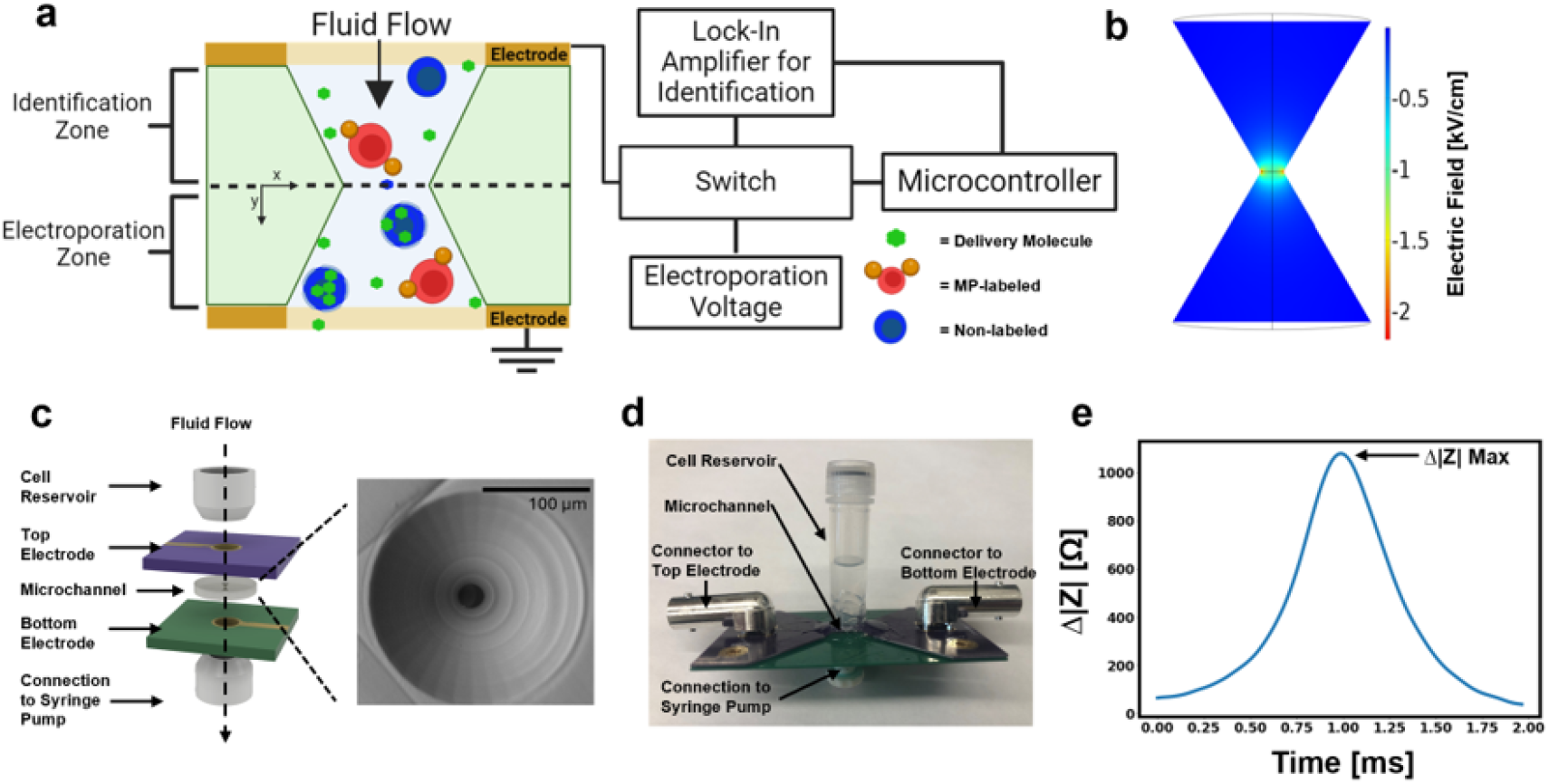
Schematic and concept of ME-SPICy: (a) Overview of ME-SPICy concept, as individual cells flow through a narrowing micro-aperture, cells are identified by a multi-parameter impedance signature measured with a lock in amplifier and analysed in real-time on a microcontroller and if a target cell is identified, the microcontroller switches the voltage applied to the top electrode (b) Simulated electric field in the micro-aperture with 15 V applied to the top electrode (c) Expanded view of the device with the 3D printed micro-aperture sandwiched between two PCB electrodes which are connected to the cell reservoir and syringe pump. SEM of the top view of the 3D printed micro-aperture is included (d) Assembled flow-cell set up (e) Impedance magnitude change (at 45kHz measurement frequency) from baseline impedance of the system for a single Jurkat cell passing through the system

As a cell flows through the first half of the constriction, sinusoidal (AC) sensing voltages are applied by a lock in amplifier (LIA) to identify the cell via real-time multifrequency impedance cytometry, wherein the whole impedance spectrum of each cell can be rapidly captured as it passes by. The LIA outputs these multiparameter impedance signals, which are high pass filtered and digitized with an analog-to-digital converter for real-time analysis by a microcontroller. The microcontroller applies an algorithm to detect maximums in impedance corresponding to the passage of individual cells. The magnitudes of the maximums across multiple frequencies are then simultaneously analyzed to discriminate MP-labelled cells, which exhibit distinct impedance signatures compared to non-labeled cells. If a target cell for EP is identified, using pre-defined signatures, as determined in real time by the microcontroller algorithm, it triggers an external switch to redirect the signal source at the electrodes from the LIA to a DC voltage supply, enabling EP before the cell leaves the aperture. A full block diagram of the electronic system can be found in Figure S1. Thus, the initial region of the microchannel is called the “identification zone” as cells are sensed until the narrowest part of the constriction, and the second half of the constriction is the “EP zone”.

Here, this system uses a micro-aperture, 3D printed in biocompatible non-conducting resin (see Methods), with the smallest dimension 25 µm in diameter, sandwiched between printed circuit board (PCB) electrodes with plated-through holes (PTHs) (Figure 1 c, d). These PCBs are then assembled with a cell reservoir and flow connector, to form a flow-cell. As the cells flow through, the rise in aperture impedance due to the cell is measured via the PCB electrodes (Figure 1e). Notably, this device does not require any complex multi-step microfabrication process to fabricate as it is directly 3-D printed from a design file enabling flexibility of aperture shape. It enables the use of low voltages (<16 V) via electric field focusing, making it much simpler, reusable, and lower cost to manufacture compared to standard PDMS channel and microelectrode-based microfluidic EP devices. Typically, PDMS-based EP devices need multiple microfabrication steps including microelectrode and mold fabrication, soft lithography and aligned bonding of PDMS on microelectrodes and yet result in devices which are often not reusable due to clogging or electrode damage via corrosion due to electrolysis.

### Simulation of Effects of MP on Cell Impedance Measurements

To enable selective in-flow EP, it is critical to identify the MP-labeled subpopulation in real time. Cell subpopulation identification is often based on surface marker expression and is typically achieved using fluorescent labeling which is detected optically. However, previous groups have shown that surface markers can also be detected electronically by labeling them with MPs [25, 29]. To assess the feasibility of electronic MP detection in our system and to guide the selection of MP size, we performed a series of simulations in COMSOL Multiphysics. We first developed a model which replicated the experimental baseline and single-cell multifrequency impedance measurements in the system (Figure S2). As shown in Figure 2a, the maximum impedance change, or where the signal to noise ratio is the highest, occurs at the location of the maximum electric field at the narrowest part of the microchannel. This indicates that peak impedance should be used for MP discrimination, and thus subsequent simulations were focused on the MP-cell interaction at this location.

**Figure 2.**
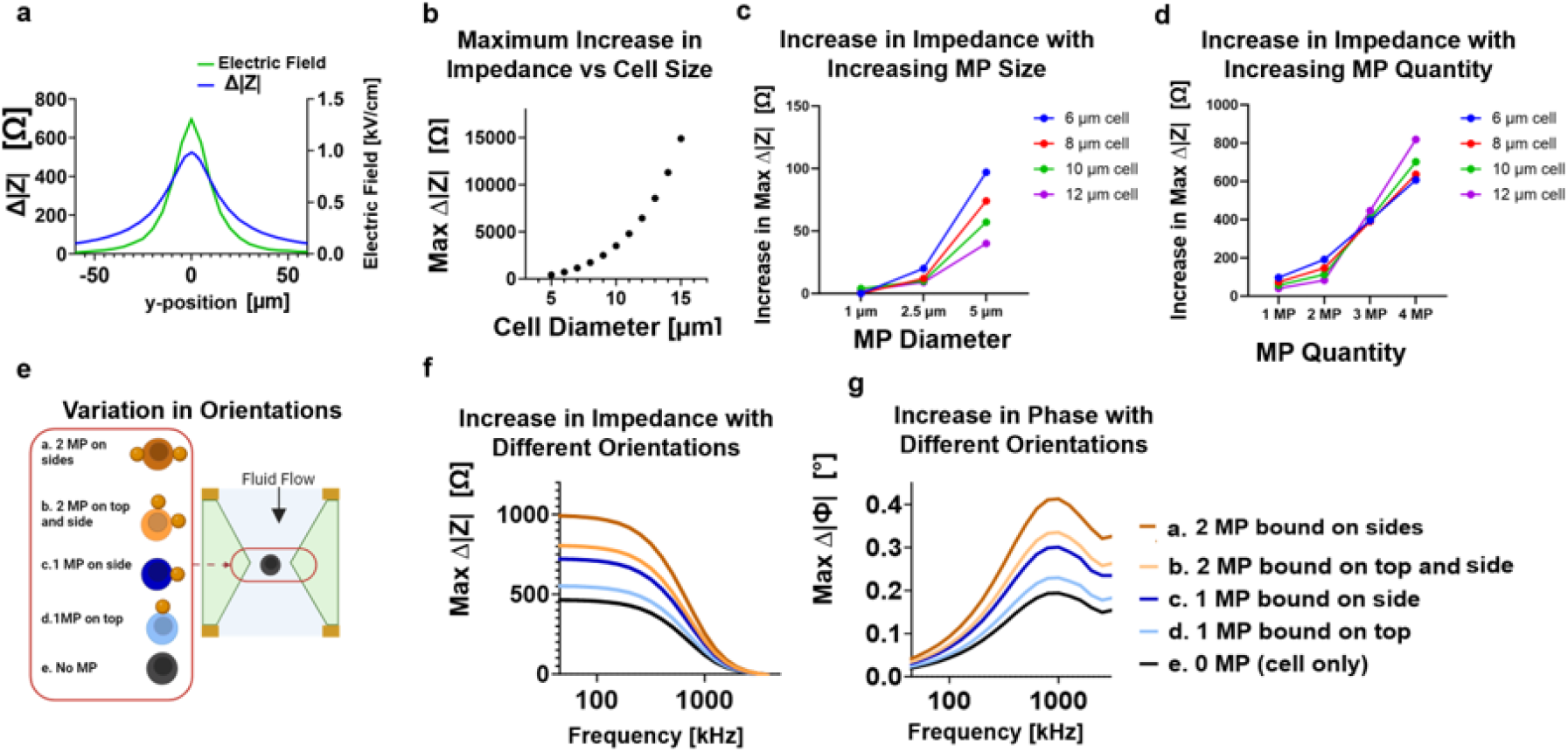
Simulated effects of MPs on the impedance of cells (a) Change in impedance of a single cell, 6 μm in diameter, with varying vertical position in the micro-aperture, where y = 0 μm is the narrowest part of the constriction. The electric field along the central axis of the microchannel vs position is overlayed. (b) Maximum impedance change from baseline detected with different cell sizes at the narrowest part of the constriction, (c) The increase in maximum impedance change of a cell caused by one MP of different diameters directly on top of the cell surface, (d) The increase in maximum impedance change of a cell caused by varying quantities of 5 μm MPs on the cell surface. Orientations of these MPs can be found in Figure S3. All impedance changes in a-d are at a frequency of 45kHz. (e) Example orientations of the MPs on a cell as it flows through the microchannel which correlate to the positions in f and g. (f,g) Impedance and phase spectra of 5 μm MP labeled cells at the different orientations shown for a cell 6 μm in diameter.

Initially, we modeled non-labeled cells as conductive spheres of different sizes, each surrounded by a thin non conducting shell representing the cell membrane. The results indicate that as the cell diameter increases, the maximum change in impedance increases at low frequencies (45 kHz) (Figure 2b). This is consistent with the Coulter principle which relates low-frequency impedance change caused by the presence of a sphere in a microchannel, or **Δ*R***, with the sphere diameter, d, and the diameter of the microchannel, D, and the resistivity of the suspension media, ***ρ***, assuming that D>>d [30]:

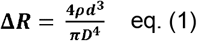

This cubic relationship shows that the volume of the sphere, or cell, drives changes in low frequency impedance. When a cell is labeled with an MP, they increase its effective volume, leading to a greater **Δ*R***. For our system, assuming the total change in impedance is additive with volume, and disregarding that D changes slightly even in the narrowest part of the micro-aperture, the total impedance change when MPs label a cell in ME-SPICy can therefore be approximated as:

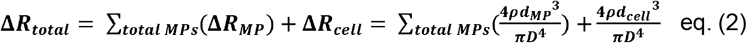

Because the impedance change scales with volume, detectability can theoretically be improved by using many small MPs or few large MPs. To evaluate this, we simulated varying diameters (1, 2.5, or 5 µm) and quantities (1-4 MPs) of MPs on cells ranging from 6-12 µm in diameter (Figure 2 c, d). To investigate MP size, a single MP was simulated directly attached to the top pole of the cell. The simulations also indicated that MPs located on the vertical poles of the cell have less of an effect on impedance compared to MPs bound the perpendicular pole of the cell. This is likely due to the proximity to the highest electric field regions along the microchannel walls (Figure 1b). Thus, to understand the minimum expected impedance change, MPs were simulated on the vertical pole, Figure 2c. These results confirm that larger diameter MPs increased the impedance more than smaller diameter MPs. However, since MPs are smaller than cells, the overall impedance change remains cell dominated. This explains why, in Figure 2c, the larger cells have a smaller increase in impedance change due to the MP constituting a smaller fraction of the total volume.

To isolate the effect of MP quantity, we simulated cells labeled with increasing numbers of the 5 µm MPs, which were identified as the most effective size in Figure 2c. Figure 2d demonstrates that impedance change increases with increasing MP quantity. MP positions used in these simulations can be found in Figure S3. Interestingly, when four MPs are present (with two on the vertical poles and two on perpendicular poles), larger cells experience the greatest impedance change, contrary to the trend in Figure 2c. This is likely due to edge effects of the electric field, because at the perpendicular poles the MPs and large cells interact more with the regions of highest electric field than small cells (Figure S3).

However, 5 µm MPs across all cell sizes and quantities cause an increase in impedance and thus were chosen for experiments. Since increasing MP quantity enhances signal, we aim to maximize the MP-cell ratio to increase the likelihood of multiple MPs binding to a single cell in experiments.

Given the position dependent edge effects observed at low frequencies, causing the approximation involved in eq. (2) to no longer be applicable as D>>d is no longer true, we also characterized different MP locations across the full frequency spectrum in the range that our system uses, Figures 2f and g. These simulations confirmed that MPs bound on the sides result in a greater change in impedance than those on the vertical poles across all frequencies. In heterogenous populations, this positional sensitivity at a single frequency could confuse the system as to whether a cell is a large cell with no MPs or a small cell with many MPs [31]. To address this, we use multiparameter impedance measurements, which have previously been used to electronically discriminate between MP-labeled and non-labeled cells [25].

As shown in Figures 2f and g, MP labeling increases the change in impedance magnitude and phase across the spectrum. Impedance is magnitude is constant at lower frequencies then falls at higher frequencies, while phase steadily increases until approximately 1 MHz This spectrum shape matches our previous work, and reflects the resistive and capacitive effects of the cell and MP [28, 31]. In impedance cytometry literature, a cell is often approximately modeled as a resistor and capacitor in series [32]. At low frequencies, the cell and MP thus act as a non-conductor impeding the flow of current which causes the volume dependent impedance change. However, at higher frequencies, ∼1 MHz [33], capacitive effects become significant, resulting in lower impedance and increase phase shift so that Eq. 1 no longer directly applies (Figure 2g). These capacitive effects can arise from the non-conducting cell membrane and MP acting as additional dielectric layers where charge can accumulate.

Together, these simulations indicate that multiparameter impedance measurements are required to accurately discriminate between labeled and non-labeled cells for selective EP. Also, these simulations show that the impedance will increase at low frequencies, which is volume driven, and at higher frequencies, which is capacitance driven. The ratio of high to low frequency impedance, also known as opacity, can likely then be a useful metric for distinguishing between non-labeled and labeled cells [25].

### Verification and Electronic Analysis of MP-Labeling of Cells

To experimentally validate and quantify MP labeling on cells, Jurkat cells were incubated with anti-CD45 conjugated 5 μm MPs and analyzed with imaging cytometry to quantify MP labeling (Figure 3a). Labeled cells exhibited higher side scatter (SSC) than non-labeled cells, with increased SSC indicating a greater number of MPs bound, (Figure 3b). Based on SSC and confirmed visually with the images from the imaging cytometer, cells were gated into groups based on the number of MPs bound. 85% of all cells had at least one MP bound, with ∼50% of cells having three or more MPs (Figure 3c). Due to the clear correlation between SSC and MP labeling, flow cytometry was used for MP labelling verification in further experiments. Flow cytometry data from this imaging cytometry experiment is included in Figure S4.

**Figure 3.**
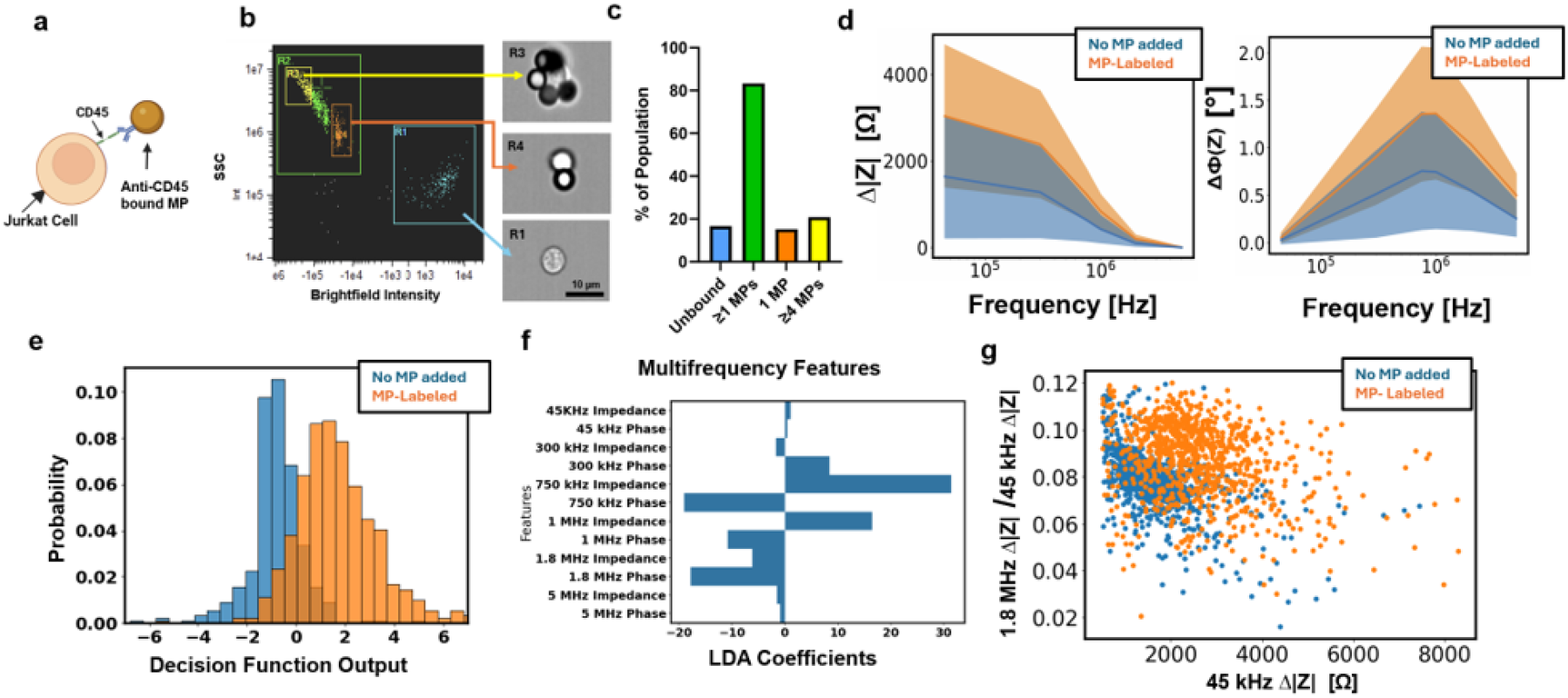
Verification of Jurkat cells labelling with anti-CD45 conjugated MPs (a) Anti-CD45 conjugated MPs bind to CD45 receptors on Jurkat cells, (b) Imaging cytometer verified SSC correlation to the number of MPs bound to cells (c) Identification of the number of MPs cells are labeled with (d) Mean impedance and phase spectra of non-labeled (control) and MP-labeled cells. The bold line is the mean while standard deviations are the highlighted regions (e) LDA separation of populations with and without MPs (f) Loadings plot of the LDA coefficients (g) Opacity plot of non labeled and labeled cells, using only 45 kHz and 1.8 MHz, control n = 1492, MP labeled n = 841.

Next, populations that were MP-labeled (Jurkats + MPs) or had no MPs added (Jurkats only) were independently flowed through ME-SPICy for multifrequency impedance characterization. This allowed simultaneous measurement of impedance and phase magnitudes at six frequencies (from 45kHz to 5MHz) at the single cell level. This data was then analyzed to find the maximum change in the impedance and phase signal for individual cells at each frequency. Population level results (n > 800) of the maximum changes are shown in Fig 3d. These results showed significantly higher (p < 0.05) impedance magnitude and phase across all frequencies for MP-labeled cells (Table S1). These findings were consistent with simulation predictions that MPs would cause an increase in impedance, driven by increased volume and increased capacitive effects. It should be noted that the population mean for the MP-labeled cells in Figure 3d, is likely an underestimate for the maximum change in impedance due to the presence of the small fraction (∼15%) of non-labeled cells in the MP-labeled population.

Linear discriminant analysis (LDA) was then applied to the dataset using the maximum impedance and phase values. Using the full feature set (12 parameters per cell – impedance magnitude and phases at 6 frequencies) achieved a separation accuracy of 0.85 (Figure 3e). The LDA coefficients, shown in Figure 3f, indicate the importance and direction of the features in separating the two populations. We see that frequencies between 750 kHz and 1.8 MHz drive discrimination between the populations, likely due to the capacitive effects of the MPs on the cell surface, which simulations showed as well. Due to better electronic signal-to-noise performance, the values at 1.8 MHz were used to plot opacity, or the ratio of high to low frequency impedance magnitude (Figure 3g). This opacity plot shows that given just the low frequencies value there is only a slight increase in impedance magnitude with the MP-labeled population (x-axis, volume driven). However, the opacity shows a clear increase for the MP-labeled population (y-axis). Because of this clearer discrimination between populations, only opacity with these two frequencies was used in further experiments.

### Selective EP of Non-labeled and MP-labeled Cells

Initially a model experiment was designed with a mixture of MP-labeled and non-labeled Jurkat cells, targeting one of the populations for EP. Here, the decision to apply the EP voltage is based on whether the single cell opacity, calculated by the microcontroller in real time, falls above or below a threshold that identifies MP-labeled Jurkats. To determine where to set the threshold, non-labeled and MP-labeled cells were first independently flowed through ME-SPICy, using the microcontroller to record the opacity calculated with 1.8 MHz and 45 kHz (Figure 4a). The numerical range of opacity values is different here than in Figure 3g due to increased amplitudes used now for the 1.8 MHZ and 45kHz signals output by the LIA. The EP threshold was then set at 95% of the opacity of the non-labeled population. Labeled and non-labeled populations were then mixed for EP to ensure both populations would be present in the final mixture of cells used for selective delivery. Unbound MPs were also present in this mixture; however, their low frequency impedance was significantly lower than that of the cells so they were electronically filtered out to avoid unnecessary switching to the EP voltage which could cause electrolysis.

**Figure 4.**
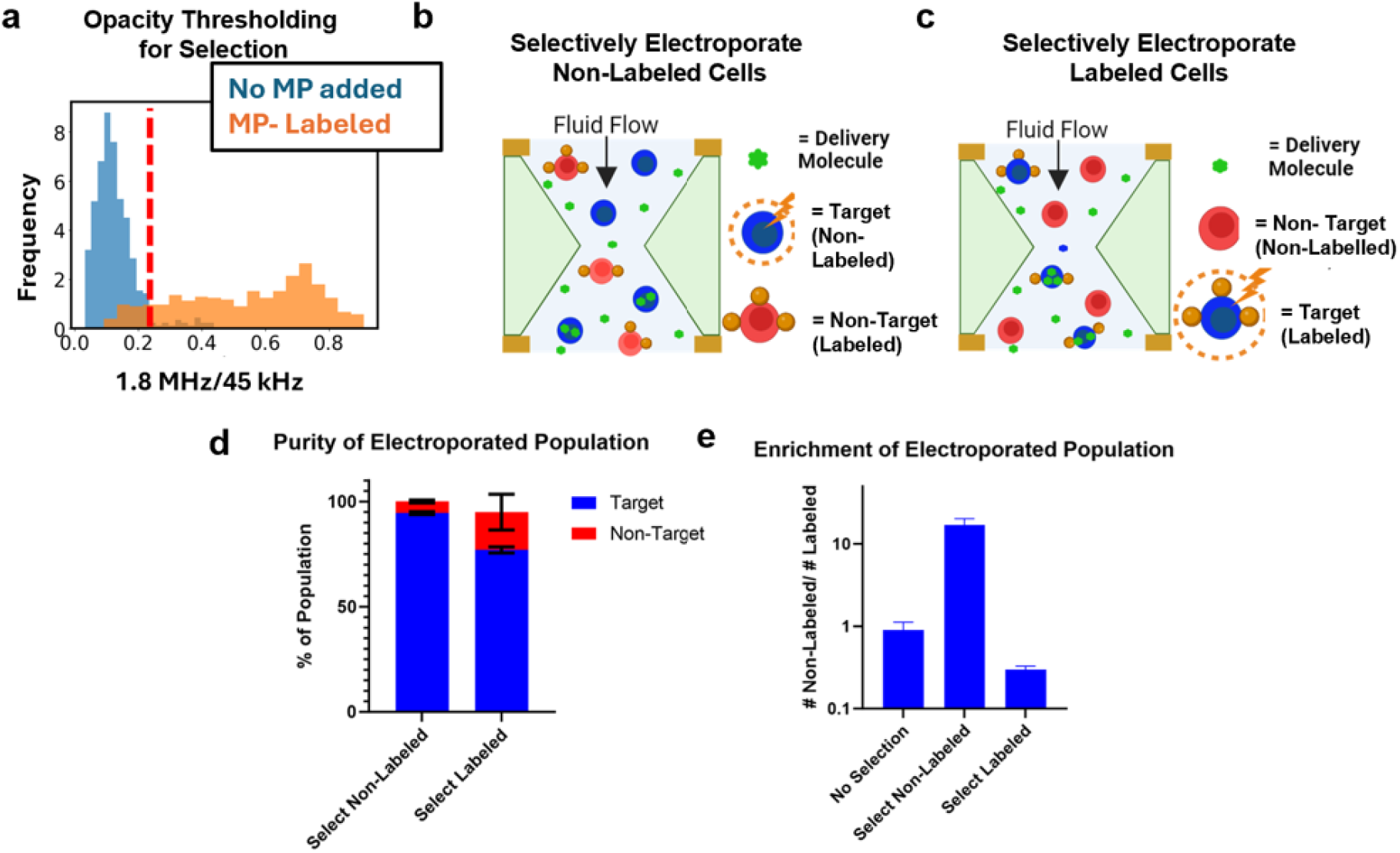
Selective electroporation of labeled and non-labeled Jurkats. (a) The multifrequency opacity threshold, red line, set for targeting either non-labeled or labeled cells, n = 1000. (b) Scheme for selecting non-labeled cells as the target cell, (c) Scheme for selecting labeled cells as the target cell, (d) Percent of the cells that were delivered to that are on target vs off target. All data are from n=2 technical repeats. Error bars shown are standard deviations, (e) Ratio of the number of non labeled cells to the number of labeled cells within the population of cells that are live and electroporated with and without selection.

Selective EP was tested by either selecting the non-labeled population, Figure 4b, or the labeled population, Figure 4c, as the target population. Propidium Iodide (PI), a fluorescent dye, was delivered and delivery efficiency was defined as the percentage of live and electroporated (PI+) cells out of all live cells. Flow cytometry was used to analyze viability, delivery efficiency, and to identify the labeled cells. For both schemes, it was found that >80% of the electroporated population was on target, with selecting for non-labeled cells yielding ∼95% purity of the target population in the electroporated population of cells (Figure 4d). Additionally, comparing the initial ratio of target to non-target cells with the final ratio of electroporated cells revealed an >10-fold enrichment of the target population when selecting non-labeled cells and ∼4-fold enrichment when targeting labeled cells (Figure. 4H). Cell viability remained high, >75% across the selection trials, and both cell viability and delivery efficiency are shown in Figure S5. Intriguingly, delivery efficiency was found to be higher when selecting for non-labeled cells than when selecting for MP-labeled cells, likely causing the lower enrichment factor when selecting labeled cells. To improve this efficiency the opacity threshold could be optimized, however this could be at the cost of purity in the electroporated sample representing a typical yield vs purity trade-off in any selection method.

### Selective EP of Lymphocytes in PBMC

Having demonstrated selective EP with a cell line, ME-SPICy was next used to selectively electroporate primary cells. Given the growing demand for lymphocyte-based cell therapies, such as NK and CAR-T cell therapies, we aimed to selectively deliver to lymphocytes in primary human peripheral blood mononuclear cells (PBMC). PBMCs contain lymphocytes (NK, B, and T cells) and monocytes, which comprised ∼38% of the PBMC batch used, as determined by flow cytometry (Figure S6).

To optimize for purity of the non-labeled fraction, monocytes were labeled via anti-CD14 conjugated MPs, resulting in a mixed population of non-labeled lymphocytes and labeled monocytes (Figure 5a). Flow cytometry confirmed that when MPs were added to PBMC, ∼80% of the monocytes are labeled, leaving the non-labeled target population to primarily consist of lymphocytes (Figure 5b). Of the cells exposed to MPs, the labeled population, on the other hand, does consists of ∼40% lymphocytes, accounting for 25% of the total lymphocyte population. While this MP labeling could be optimized further, via changing the antibody concentration, the bead-to-cell ratio, or the kind of MP used, the 90% purity of lymphocytes in the non-labeled population was considered sufficient to proceed with selective EP of non-labeled cells.

**Figure 5.**
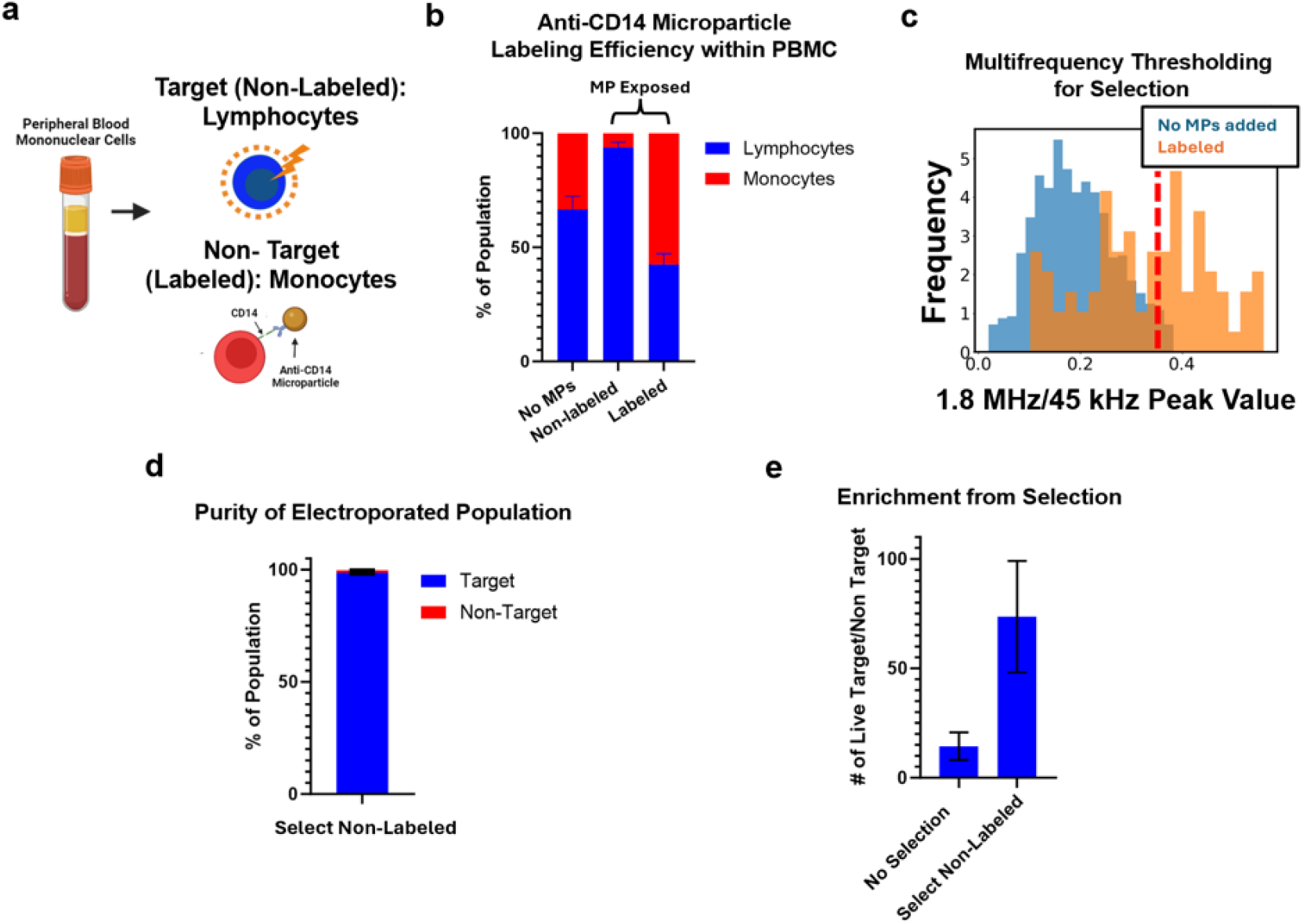
Selective electroporation of non-labelled lymphocytes from PBMC (a) PBMC are labeled with anti-CD14 conjugated MPs. Non-labeled lymphocytes are selected as the target for EP (b) Proportions of lymphocytes and monocytes in the labeled and non-labeled populations, (c) The multifrequency opacity threshold, red line, set for targeting non-labeled lymphocytes, (d) Percent of the cells that were delivered to that are on target vs off target. All data are from n=2 technical repeats. Error bars shown are standard deviations, (e) Ratio of the number of non labeled (target) cells to the number of labeled cells within the population of cells that are live and were electroporated with and without selection.

As was observed previously, there is a clear shift in the opacity of MP-labeled primary cells (Figure 5c), with a high opacity population emerging with the addition of MPs similar to what was seen with MP-labeled Jurkat cells. The full frequency spectrum of the measured impedances can be found in Figure S7. Thus, a threshold for selection to include 95% of non-labeled PBMC (Figure 5c), was determined. Below this threshold, non-labeled cells, which are primarily lymphocytes, were selectively electroporated, while cells above the threshold were identified as MP-labeled and were not electroporated.

PI was delivered to the non-labeled lymphocyte population. Similarly to the Jurkats, a purity of ∼95% was achieved in the electroporated cells, meaning that 95% of the cells that were electroporated were on target, Figure 5d. This enabled the ratio of electroporated target to non-target cells to increase by factor of (Figure 5e) enriching the non-labeled population. Delivery efficiency was similar to that achieved with Jurkat cells. However, cell viability was found to be slightly lower compared to with the Jurkat cells, which can be due to the largest proportion of cells in this non-labeled population being non-activated T cells, which can be difficult to transfect and can suffer from lower viability upon electroporation [34]. Given the wide parameter space for the EP voltage, such as number of pulses, their length, and amplitude, there is considerable room for further optimization to improve both delivery efficiency and viability.

A known limitation of many single-cell microfluidic techniques is their throughput. Although ME-SPICy can currently process up to 0.5-1 million cells per hour, this remains lower than other bulk or macro-fluidic EP approaches [13, 35]. Throughput could be increased by decreasing the time spent in the identification zone and electroporation zones. Decreasing identification time could be accomplished by using a faster flow rate, higher cell concentrations, or different identification algorithms. Additionally decreasing the EP pulse time or parallelizing with multiple microchannels would allow for significant increases in throughput. Here, a significant advantage of the ME-SPICy approach is that it is inherently easier to scale due to the flexibility provided by the way it is designed and fabricated, as described earlier.

With further optimization of MP labeling, ME-SPICy could also be used to target specific lymphocyte subpopulations such as NK cells, CD8^+^ cytotoxic T cells, or CD4^+^ helper T cells. Beyond lymphocytes, this strategy could also be adapted for stem cells or other clinically relevant rare populations. Although current selection relies on MP labeling, future integration of cleavable antibodies, shear-removable MPs, or magnetic separation could facilitate MP removal post-electroporation [36].

We also note that while EP coupled with downstream label-based sorting can accomplish similar selection goals, ME-SPICy simplifies such workflows and enables additional functionalities through real-time single-cell impedance measurement. For example, the platform could be adapted for selective killing, applying high-voltage irreversible EP to eliminate unwanted cells, such as cancerous B cells in CAR-T manufacturing or even circulating tumor cells (CTCs) [37]. Furthermore, the impedance-based feedback control in the system could enable dynamic EP voltage modulation, adjusting pulse strength based on cell size or other properties, reaching a threshold that relates to permeability, thereby improving delivery precision [19].

## Conclusions

We developed a microfluidic EP platform capable of achieving high-purity selective EP of labeled subpopulations and demonstrated it with both immortalized cell lines and human primary cells. ME-SPICy adds a new dimension to EP which at the commonly used bulk scale, lacks selection or targeting ability. It has thus the potential to eliminate the need for cell sorting for workflows that use EP such as cell engineering, reducing costs from reagents and equipment, and simplifying the cell manufacturing process. Moreover, the performance of ME-SPICy is on par with conventional bulk EP in terms of delivery efficiency, with potential for further optimization to increase efficiency and throughput, broadening its clinical applicability. This microfluidic platform also allows for flexibility and integration into existing manufacturing workflows, offering an alternative to traditional viral methods with additional benefits for integration. This positions ME-SPICy as a transformative tool for both research and clinical applications in cell engineering. **Experimental Methods**

### Finite-Element Modeling and Simulations

Finite element analysis was performed in COMSOL Multiphysics 5.6 using the AC/DC and electrical circuit modules. The electrical circuit module was used to couple circuit elements to match the baseline experimental impedance of the device. Cells were modeled as spheres of conductive media, σ = 1.1 S/m, using the contact impedance feature to model the cell membrane as a thin non conducting layer. Microparticles were modeled as solid non conducting spheres.

### Device Fabrication

ME-SPICy consists of a narrowing vertical micro-aperture which is 3D-printed using a Nanoscribe Photonic Professional GT2 and sandwiched, using epoxy, between 2 PCBs with gold-coated copper electrodes. Further details of can be found in previous work [28, 31].

### Cell Culture

Jurkat cells (Clone E6-1, Fisher Scientific) were cultured in RPMI-1640 medium (Thermofisher, A1049101) supplemented with 10% Fetal Bovine Serum (FBS) and 1% Penicillin/Streptomycin. Cells were passaged every 3-4 days and kept in a humidified incubator at 37°C. Human PBMC, purchased from STEMCELL Technologies (70025), were aliquoted, re-frozen, and kept in liquid nitrogen. For experiments, Jurkat cells, from culture, or PBMC, after thawing for 10 minutes, were washed with PBS for 5 minutes at 400 g. Jurkat cells were then resuspended at 3×10^6^ cells/mL of delivery buffer (90% Opti-MEM (Thermofisher, 31985070), 10% OptiPrep (STEMCELL Technologies, 07820)). For microparticle labeling, Jurkat cells were suspended in PBS with 1% BSA instead. PBMC were resuspended at 3×10^6^ cells/mL in PBS 1% BSA and then were stained with CD3, CD19, and CD56 for 30 minutes at room temperature before being washed in PBS with 1% BSA for verification of cell subpopulations. They were then washed and resuspended at 3×10^6^ cells/mL in PBS with 1% BSA for labeling.

### Microparticle Labeling

5.7 μm polystyrene streptavidin functionalized MPs were obtained from Spherotech (SVP-50). MPs were vortexed and incubated for 30 minutes at room temperature on a shaker with anti-CD45 biotin (ThermoFisher, 13-0459-82) for labeling Jurkats, or with anti-CD14 for PBMC (ThermoFisher, 13-0149-82) for labeling monocytes. 2.5 μg of antibody was added for 1 x 10^5^ beads. The MPs were then washed with PBS with 1% BSA once at 3000g for 3 minutes before adding cells suspended in PBS with 1% BSA at a concentration of 3×10^6^ cells/mL, at a ratio of ∼40 MPs per cell. The cell and MP mixture was then incubated for 30 minutes at room temperature on a shaker. For impedance characterization the cells were left in PBS, but for EP experiments the mixture was diluted in delivery buffer. Verification of MP binding was achieved by using an Amnis MKII imaging cytometer and a Cytek Aurora Flow Cytometer.

### Impedance Measurements for Machine Learning

A lock-in-amplifier and transimpedance amplifier (LIA) (HF2LI, HF2TA, Zurich Instruments) were used to detect cells as they flow through the aperture, only interfacing ME-SPICy to the LIA with no additional circuit elements. Impedance and phase were recorded at 6 frequencies, ranging from 45 kHz to 5 MHz at 1 V each. Cells were pulled through the system at 1 μL/minute via a syringe pump, Chemyx f200x. A peak finding algorithm is then applied to this data to identify the cell peaks at the lowest frequency, filtering out noise and unbound MP. The time index at which this peak is identified is then used to extract the remaining change in impedance and phase measurements at other frequencies. Linear discriminant analysis (LDA) was then used to reduce the dimensionality of the data to identify the two most relevant frequencies that would be used for EP experiments. The EP circuit is external to the LIA but uses the LIA auxiliary outputs which relate to 1 selected frequency each. The training data for the LDA model was datasets of the peak impedance and phase values at the different frequencies for non-labeled cells and MP-labeled cells. The data was standardized and split so that 70% was used for training and 30% for testing.

### Selective EP Control Mechanism

A 3V 45 kHz and a 4.5V 1.8 MHz sine wave were applied to the top PCB electrode by the LIA for impedance detection. The lock-in amplifier then internally demodulates and amplifies the signal 300 times and removes the DC offset. The signals from the individual frequencies are then output by the LIA, through a 1^st^ order high pass filter (f_c_ = 70 Hz), to an analog to digital converter, AD7366 (Analog Devices), which communicates to a Teensy 4.1 microcontroller at 160,000 samples per second. The microcontroller is constantly running a peak finding algorithm on the moving average of the signal to detect cells in the micro-aperture, filtering out noise and any MPs not attached to cells. Once a cell peak has been identified, if the ratio of high to low frequency impedance (opacity) is within a set target range, the microcontroller sends a signal to an external switch (TMUX8109) which switches the voltage applied to the top electrode from the detection voltage to a 15.5 V DC EP voltage for 1 ms before switching back to the detection voltage. If a cell is not within the target opacity range, there is no switching.

### Selective EP

PBS with 1% BSA was flowed through the aperture prior to beginning experiments for 15 minutes. The impedance data of unbound cells and labeled cells in delivery buffer were individually recorded before beginning EP experiments to determine where the opacity range should be set for selective EP of the target cells. The bounds of this range were set to include 95% of the target population. For Jurkat cell EP, non-labeled and labeled Jurkat cells were mixed at a ratio of 1:2 since most Jurkats exposed to MPs were labeled and experiments needed to include a non-labeled cell population. For PBMC EP, the lymphocyte population was the non-labeled population. For each trial, cells were diluted to 1.5 ×10^6^ cells/mL with delivery buffer that contained 130 ug/mL propidium iodide (ThermoFisher, P1304MP) for a final PI concentration of 65 μg/mL. 50 μL of cells were then pipetted into the reservoir and flowed through the system at 0.5 μL/minute for 20 minutes. Post EP, cells were immediately removed from the system, stained with 1:50 volume ratio of 50 μM Calcein Blue AM for viability, and analyzed on a Aurora Cytek Flow Cytometer. Immunofluorescent staining was used to confirm cell subpopulations, and all data was analyzed in FlowJo and visualized in GraphPad Prism. PI+ classification set above the fluorescence of 95% of cells that were incubated with PI but were not electroporated. Cell viability was calculated as the number of live cells normalized to non-electroporated controls, accounting for pre-EP viability that was ∼80%.

## Author contributions

M.H. performed experiments and electronics and flow system development. J.R performed micro-aperture design and fabrication, electronics and flow system development. Y.R. performed cell culture and SEM imaging. A.S. conceptualized the project and supervised all aspects. M.H. and A.S. wrote the manuscript with inputs from all authors. All authors reviewed and approved the manuscript.

