## Supplemental for "Microparticle-Enabled Single Cell Multiparameter Electronic Immunophenotyping for Selective Electroporation"

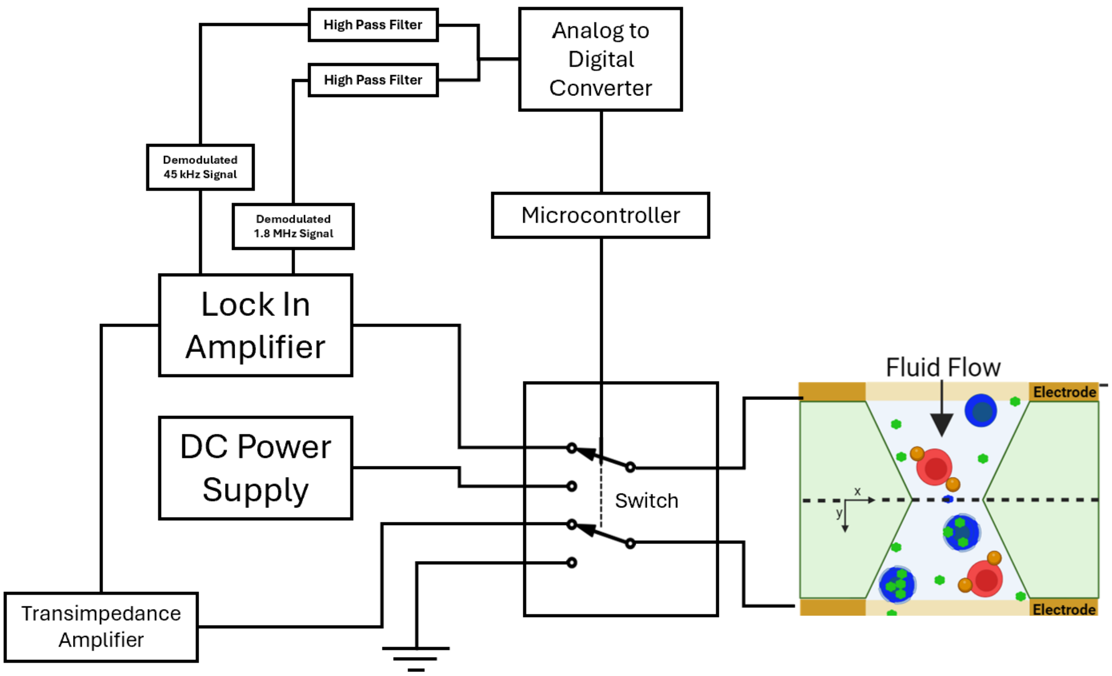


Figure S1: Block diagram of ME-SPICy. A lock in amplifier (LIA) outputs a signal to the top electrode of ME-SPICy. A transimpedance amplifier then converts the current from the system to a voltage which is then demodulated by the lock in amplifier to separate the 45 kHz and 1.8 MHz signals. These are then separately output by the LIA and high pass filtered (cutoff frequency = 70 Hz) and input to an analog-to-digital converted to digitize these signals to be read by a microcontroller. This microcontroller then analyzes this signal to identify cells and controls electroporation via a switch.

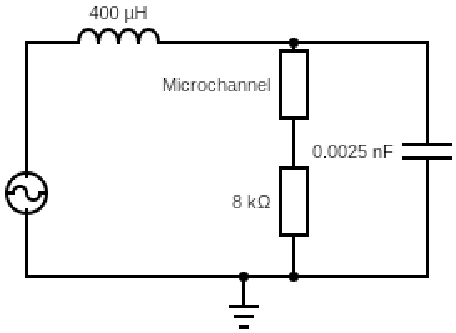


a)

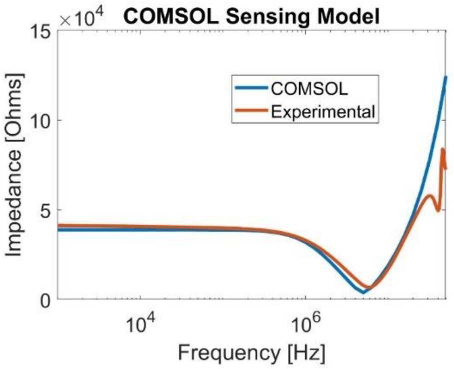


b)

Figure S2: (a) Circuit coupled to micro-aperture geometry for finite element analysis model. (b) Simulated impedance spectrum matched to experimental impedance spectrum with the microchannel filled with conducting media.

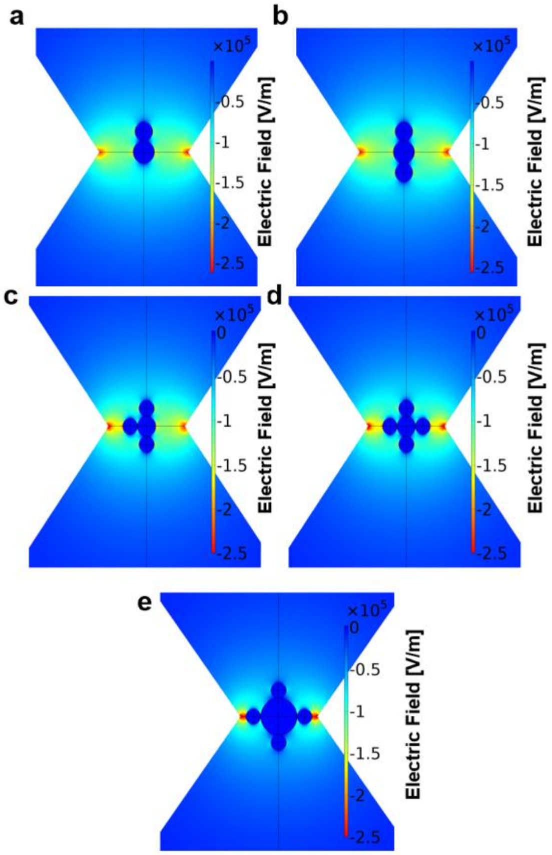


Figure S3: Positions of MPs on cells, corresponding to impedance data in Figure 2D. a) A single 5 μm MP on a 6 μm in diameter cell. b) Two 5 μm MPs on a 6 μm in diameter cell. c) Three 5 μm MPs on a 6 μm in diameter cell. d) Four 5 μm MPs on a 6 μm in diameter cell. e) Four 5 μm MPs on a 12 μm in diameter cell.

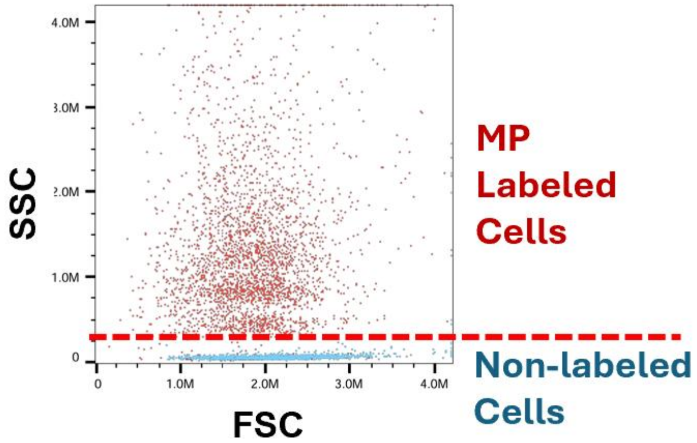


Figure S4: Flow cytometer data of MP labeled cells (red) compared to non-labeled cells which were never exposed to MPs. These cells were from the same sample that was used for imaging cytometry.

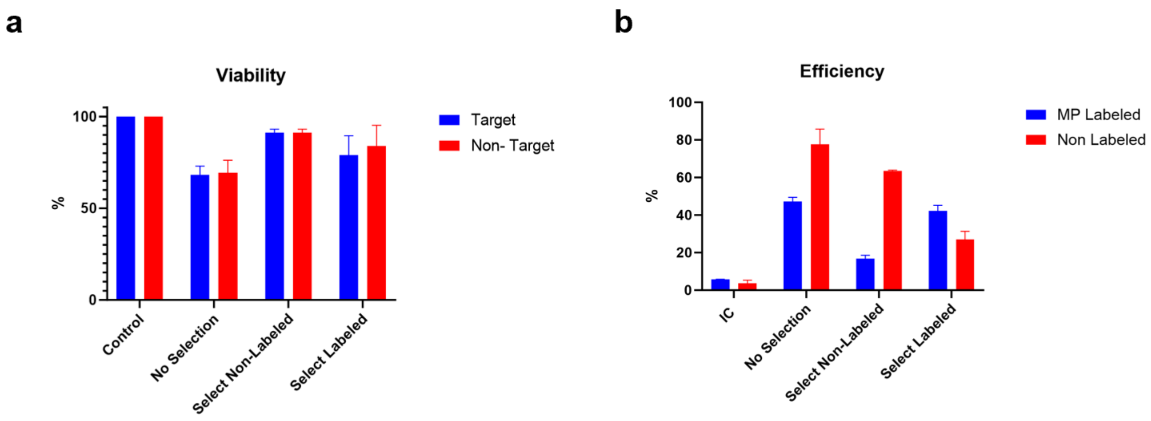


Figure S5: a) Viability of Jurkat cells, normalized to controls which were not electroporated but exposed to PI. b) Delivery efficiency of PI across the trials. N=2

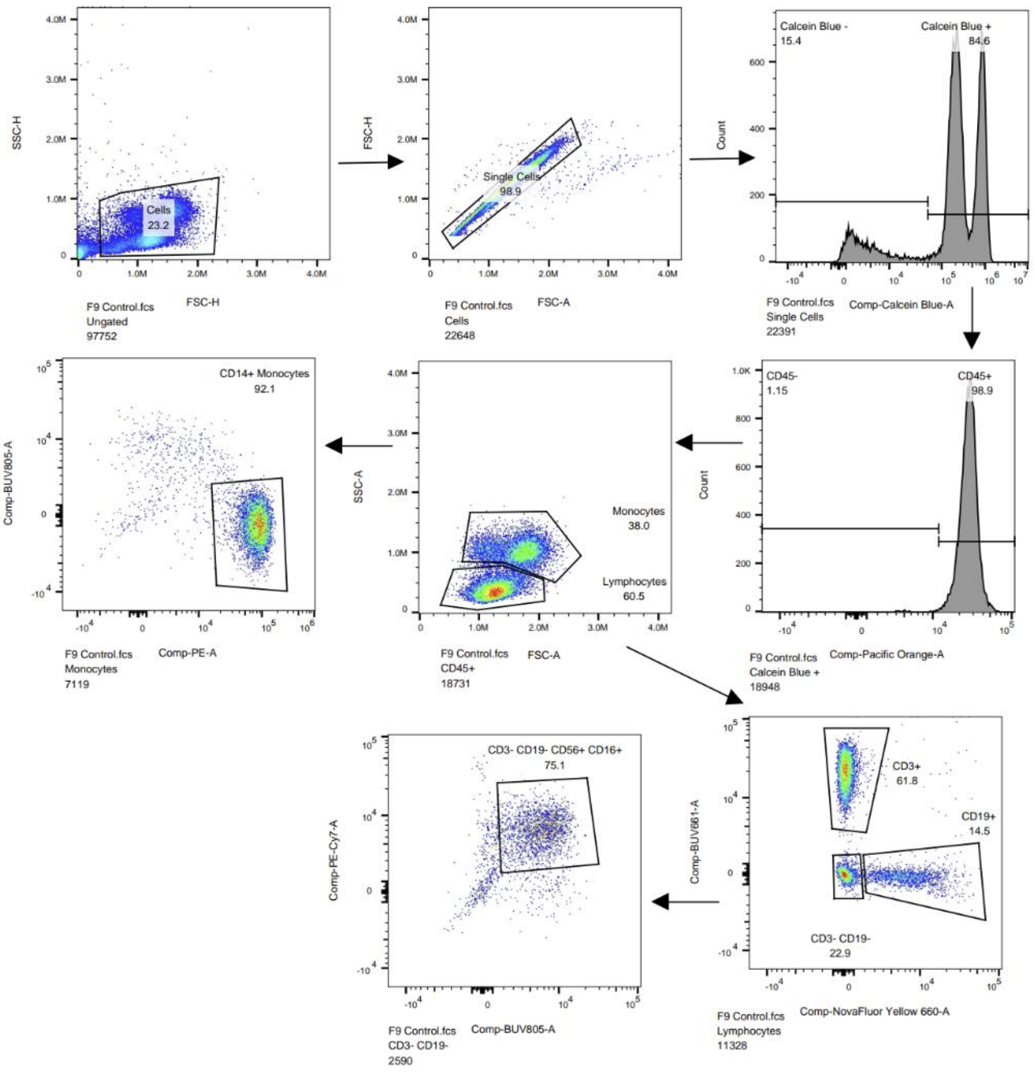


Figure S6: Antibody panel to determine the number of CD14+ cells. Cells were gated on singlets, then Calcein Blue was used for live dead staining. Live cells were that were CD45+ via pacific orange were then split into monocytes and lymphocytes via FSC and SSC. Monocytes identified by size were verified with PE which stained for CD14 and lymphocytes were stained with BUV661 for CD3, Nova Flour Yellow for CD19 for T and B cells respectively. CD3- CD19- cells were then analyzed for PE-CY7 for CD56, and BUV805 for CD16 to identify NK cells.

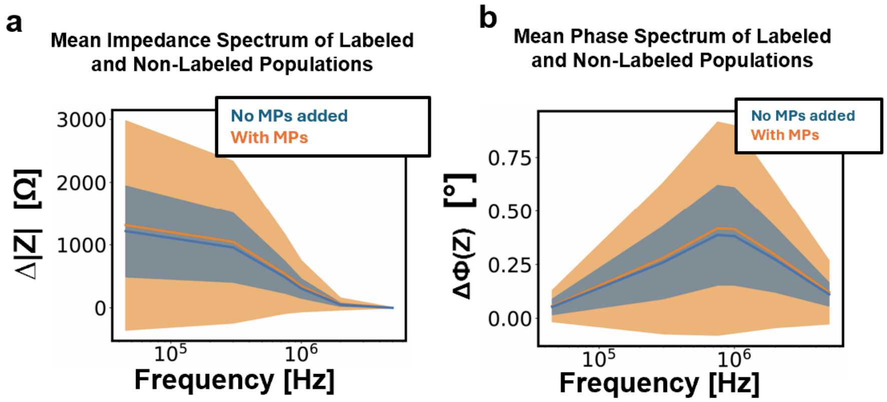


Figure S7: Impedance and phase frequency spectrum for labeled and non-labeled PBMC. Means are plotted with standard deviations plotted as the highlighted regions. The with MP population includes non-labeled lymphocytes and monocytes as well as any labeled cell.

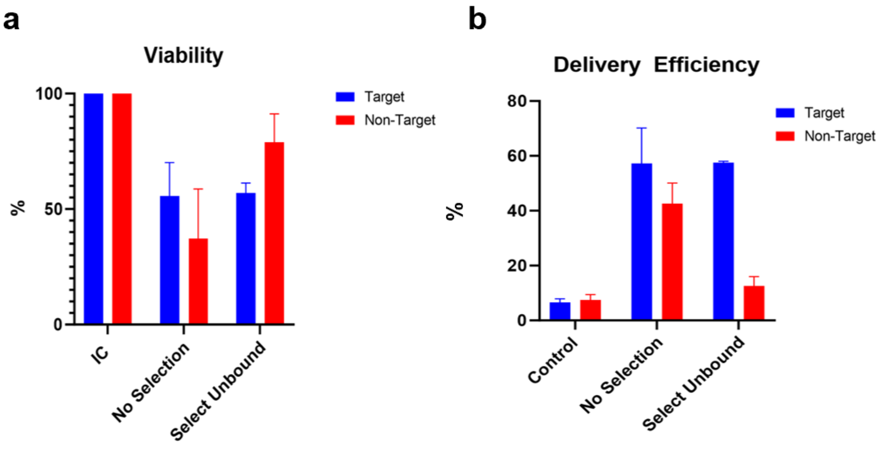


Table S1: Comparison between unlabeled and labelled Jurkat cells (t test), conducted for the impedance and phase at every frequency for n=841 cells each.

| Frequency [kHz] | Impedance p-value | Phase p-value |
|---|---|---|
| 45 | 8.82E-15 | 2.37E-05 |
| 350 | 1.41E-16 | 1.49E-11 |
| 750 | 2.73E-20 | 8.95E-15 |
| 1 | 5.25E-22 | 5.17E-16 |
| 1.8 | 7.24E-26 | 4.94E-19 |
| 5 | 9.62E-15 | 3.00E-21 |
